# Maternal SMCHD1 regulates *Hox* gene expression and patterning in the mouse embryo

**DOI:** 10.1101/2021.09.08.459528

**Authors:** Natalia Benetti, Quentin Gouil, Andres Tapia del Fierro, Tamara Beck, Kelsey Breslin, Andrew Keniry, Edwina McGlinn, Marnie E. Blewitt

## Abstract

Parents transmit genetic and epigenetic information to their offspring. Maternal effect genes regulate the offspring epigenome to ensure normal development. Here we report that the epigenetic regulator SMCHD1 has a maternal effect on *Hox* gene expression and skeletal patterning. Maternal SMCHD1, present in the oocyte and preimplantation embryo, prevents precocious activation of *Hox* genes postimplantation. Without maternal SMCHD1, highly penetrant posterior homeotic transformations occur in the embryo. *Hox* genes are decorated with Polycomb marks H2AK119ub and H3K27me3 from the oocyte throughout early embryonic development; however, loss of maternal SMCHD1 does not alter these marks. Therefore, we propose maternal SMCHD1 acts downstream of Polycomb marks to establish a chromatin state necessary for persistent epigenetic silencing and appropriate *Hox* gene expression later in the developing embryo. This is a striking role for maternal SMCHD1 in long-lived epigenetic effects impacting offspring phenotype.

## Introduction

It is now clear that epigenetic information can be passed from generation to generation via the germline, changes in which can have long-lasting effects in the offspring. One of the most notable of these effects is transmission of epigenetic information from the oocyte to the zygote. The oocyte supplies the entire cytoplasm containing all expressed mRNA and proteins to the zygote, sustaining it through its initial cell divisions until its own zygotic genome is transcribed, at embryonic day (E) 2.0 in mice ^1^. Genes whose expression is required in the oocyte for normal development of the offspring are known as maternal effect genes.

A classic example of the role of maternal effect genes in passing long-lived epigenetic information from parent to offspring is genomic imprinting, where genes are monoallelically expressed in a parent-of-origin-specific manner. Epigenetic imprints are imparted by germ cell-derived DNA methylation or trimethylation of lysine 7 on histone 3 (H3K27me3) ^2, 3, 4^. Maternal effect genes important for imprinting generally have a role in establishing and maintaining these germline marks ^5, 6^.

*Structural maintenance of chromosomes hinge domain containing 1* (*Smchd1*) is a recently defined maternal effect gene that is expressed in the oocyte and is required for genomic imprinting in the mouse placenta ^7, 8, 9^. In its zygotic form, SMCHD1 plays a key role in epigenetic silencing of imprinted loci, along with other clustered gene families and the inactive X chromosome ^10, 11, 12, 13, 14, 15^. Heterozygous variants in *SMCHD1* are also associated with the human diseases Facioscapulohumeral muscular dystrophy (FSHD) and Bosma arhinia microphthalmia (BAMS) ^16, 17, 18, 19^, demonstrating the important role SMCHD1 plays in normal development.

SMCHD1 is a member of the SMC family of proteins, large chromosomal ATPases important for chromosome structure ^20^. SMCHD1 also plays a role in chromatin architecture, mediating long-range interactions at its targets ^12, 21, 22, 23^. Recruitment to at least one of its targets, the inactive X chromosome, is dependent on the polycomb repressive complex 1 (PRC1) mark ubiquitination of lysine 119 of histone H2A (H2AK119ub) ^22, 24^. For imprinted genes we have proposed that SMCHD1 is recruited downstream of PRC2’s mark H3K27me3 ^7^. Precisely how zygotic or maternal SMCHD1 enables gene silencing is not yet clear.

One of the clustered gene families zygotic SMCHD1 binds and silences is the *Hox* genes ^11, 12, 25^, a highly conserved set of transcription factors that are responsible for correct patterning of body segments along the anterior-posterior (A-P) axis during embryonic development ^26, 27, 28, 29^. *Hox* genes are only expressed at specific times and in specific tissues during post-implantation embryonic development ^30, 31^. At all other times they are silent and marked by H2AK119ub and H3K27me3 ^32, 33^, including in the oocyte and through pre-implantation development ^34, 35, 36^, opening the exciting possibility of maternal effects on *Hox* gene expression. Based on these data and *Smchd1*’s role as a maternal effect gene, we investigated whether maternal SMCHD1 has long-lasting effects at its targets in the embryo, specifically on the *Hox* genes. This was made possible because, unlike many maternal effect genes, deletion of maternal *Smchd1* does not result in embryonic lethality ^7^.

In this study we show that maternal SMCHD1, found in the preimplantation embryo, is required to prevent premature *Hox* gene activation in the early post-implantation embryo. Interestingly, these changes occurred without disruption of H2AK119ub or H3K27me3 marks over *Hox* genes in the pluripotent state, suggesting that maternal SMCHD1 acts downstream of Polycomb to regulate *Hox* gene expression and normal skeletal patterning post-implantation.

## Results

### Maternal SMCHD1 is required for normal skeletal patterning

Given that previous work in our lab has shown that *Smchd1* mutants exhibit homeotic transformations ^12, 25^, we first assessed whether *Smchd1* maternal knockout embryos also show abnormal skeletal patterning. We set up F1 crosses between C57BL/6 and Castaneus strain (Cast) parents, using MMTV-Cre or Zp3-Cre to knock out *Smchd1* in the oocyte (Fig. 1a-c) as we have previously ^7^. We set up three types of F1 crosses. The first was a control cross yielding embryos with wild-type SMCHD1 function (*Smchd1*^wt^). This established a baseline of skeletal patterning in the F1 embryos (Fig. 1a). Almost all of these embryos had normal skeletal patterning, with 97% and 86% of control mice from the MMTV-Cre and Zp3-Cre colonies respectively having the expected 7 cervical vertebrae, 13 thoracic vertebrae, 6 lumbar vertebrae and 4 sacral vertebrae (Fig. 1d). In the second cross, *Smchd1* was deleted in the oocyte with either MMTV-Cre or Zp3-Cre, yielding *Smchd1* heterozygous embryos which lacked maternal SMCHD1 (*Smchd1^matΔ^*, Fig. 1b). The third cross, performed with the MMTV-Cre only, was reciprocal to the maternal deletion cross and generated both *Smchd1*^*del*/+^(*Smchd1^het^*) and *Smchd1^wt^ embryos*, with the oocytes from which they were generated having wild-type levels of SMCHD1 (Fig. 1c). This latter cross tested whether any phenotype observed in the *Smchd1^matΔ^* embryos was due to haploinsufficiency for SMCHD1 after zygotic genome activation rather than lack of maternal SMCHD1, and controlled for the direction of the interstrain cross.

**Figure 1.**
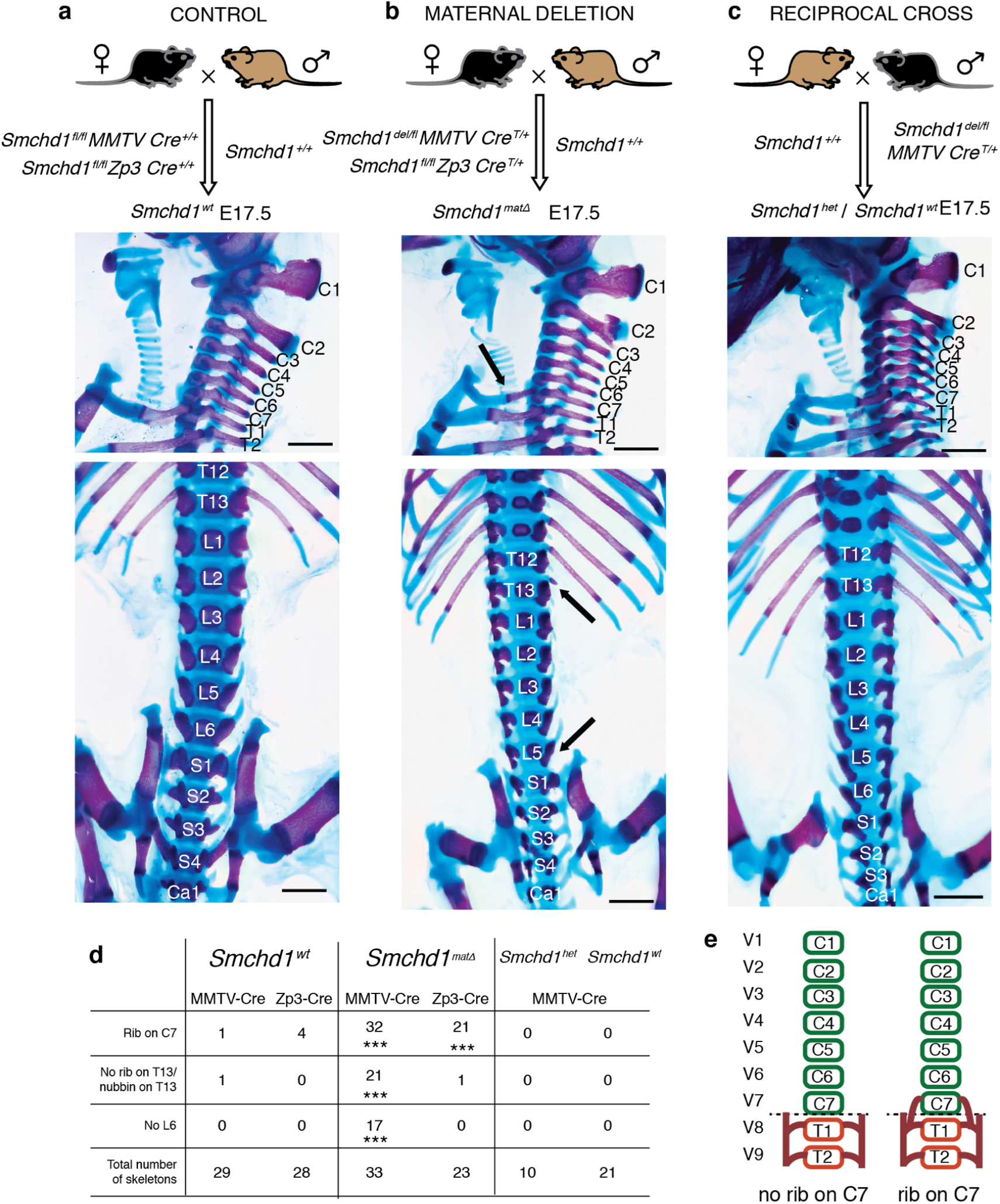
Maternal SMCHD1 is required for normal skeletal patterning. (a-c) Upper panels: the genetic crosses used to create control (a), maternal null (b) and reciprocal cross heterozygous control (c) embryos for skeletal analysis. The black mouse represents C57BL/6 strain, the brown mouse Cast strain. Middle and lower panels: E17.5 skeletons stained with alizarin red (bone) and alcian blue (cartilage), showing a sagittal view of the cervico-thoracic region and dorsal view of the thoracolumbar-sacral region. The asterisks in (b) indicate abnormalities compared with the standard axial formulae found in the controls. The skeletons shown are from MMTV-Cre crosses. Scale bar = 5 mm. (d) The summarised data for all skeletons are provided for MMTV and Zp3-Cre. The asterisks indicate statistical significance in comparing to control (chi-square test, *** p<0.001). (e) Cartoon depiction of normal skeletal patterning and the *Smchd1^matΔ^* phenotype are shown in (e) for the cervico-thoracic region.

*Smchd1^matΔ^* embryos exhibited a highly penetrant addition of a rib on the seventh cervical element (C7), suggesting that C7 adopts the identity of T1 (Fig. 1b, e). We observed several morphological variations of this additional rib including a short ectopic rib, a rib which fused with T1 with and without subsequent bifurcation before joining the sternum, and a full rib which joined the sternum independently of T1 (Supplementary Dataset 1). Grouped together, any indication of C7 transformation was observed in the MMTV-Cre and Zp3-Cre models at a penetrance of 97% and 91% respectively (Fig. 1d). Additional posteriorising transformations were observed in a subset of *Smchd1^matΔ^* embryos when *Smchd1* was deleted with MMTV-Cre. These included (*i*) the loss of ribs on T13 leading to a complete T13-to-L1 transformation or severely hypomorphic rib(s) on T13, and (*ii*) a L6-to-S1 transformation. These phenotypes were observed at a lower penetrance than the additional C7 rib; 63% and 52% for the transformation altering T13 and L6 respectively (Fig. 1d). All three of these phenotypes had significantly higher penetrance following maternal deletion of *Smchd1* compared to both the control and reciprocal cross (p<0.001, Chi-square test), and there was no sex-specificity in the phenotypes observed (Supplementary Dataset 1). Of note, when a lumbar transformation was observed, it was almost exclusively coincident with transformations at cervico-thoracic and thoraco-lumbar transitions. This suggests serial homeotic transformation in these embryos, supported further by examples of transformation of vertebra surrounding these transition points where specific morphology can be delineated (e.g. C5→C6 and T1→T2; Supplementary Dataset 1). Taken together, deletion of maternal Smchd1 results in a highly penetrant posterior homeotic transformations that can encompass multiple axial regions, implying a potential global shift in patterning effectors. Given there were no abnormalities observed in the Smchd1 heterozygous skeletons (Fig. 1c, d), these data support the view that maternal SMCHD1 is required for appropriate axial patterning.

### Maternal *Smchd1* deletion leads to precocious activation of anterior *Hox* genes

Next we investigated whether there were changes in *Hox* gene expression in *Smchd1^matΔ^* embryos that may explain the homeotic transformation phenotype. We first re-analysed our published RNA-sequencing (RNA-seq) data from control and *Smchd1* maternal null E2.75 morula ^7^. *Hox* genes were not readily detectable, and there was no change observed in the maternal null compared with control morulae for the single detectable *Hox* gene (*Hoxb13*, Supplementary Figure 1). Given the challenge with detecting low level expression in low input samples, we went on to examine post-implantation developmental stages. The tissue we chose for RNA-sequencing was tailbud tissue of the E8.0-E8.5 embryo, dissected just anterior to the level of the node. This tissue contains the caudal end of the presomitic mesoderm (PSM) and the region harbouring progenitors of the vertebral column, the neuromesodermal progenitors (NMPs) ^37, 38^. It is the *Hox* expression signatures within cells prior to somite formation that are known to instruct vertebral morphology later in development ^39^.

We conducted RNA-seq in tailbud tissue from *Smchd1^wt^* and *Smchd1^matΔ^* embryos, with somite-matched replicates (Supplementary Figure 1, Fig. 2a, b). We chose 6-11 somites as we theorised that loss of maternal SMCHD1 may lead to precocious *Hox* gene activation. There was very limited differential expression genome-wide in these somite-matched samples. Indeed, what limited differential expression was present can be explained by the sex disparity in samples at each somite number (Supplementary Dataset 2). Moreover, there was no difference in somite range between control and *Smchd1* maternal null embryos (Supplementary Dataset 2), suggesting there was no striking developmental delay following loss of maternal SMCHD1. Consistent with this, *Hox* gene expression was approximately normal in control and *Smchd1^matΔ^* tailbud samples (Fig. 2c, d). When we compared these two somite series, we saw a collective, albeit modest, upregulation of anterior *Hox* genes in somite 8-11 tissue, specifically a trend towards precocious activation of the *Hox2* to *7* paralogues in somite 10 and 11 tissue (Fig. 2e). We also observed a concomitant downregulation of posterior *Hox* genes particularly at the earlier somite stages (Fig. 2d).

**Figure 2.**
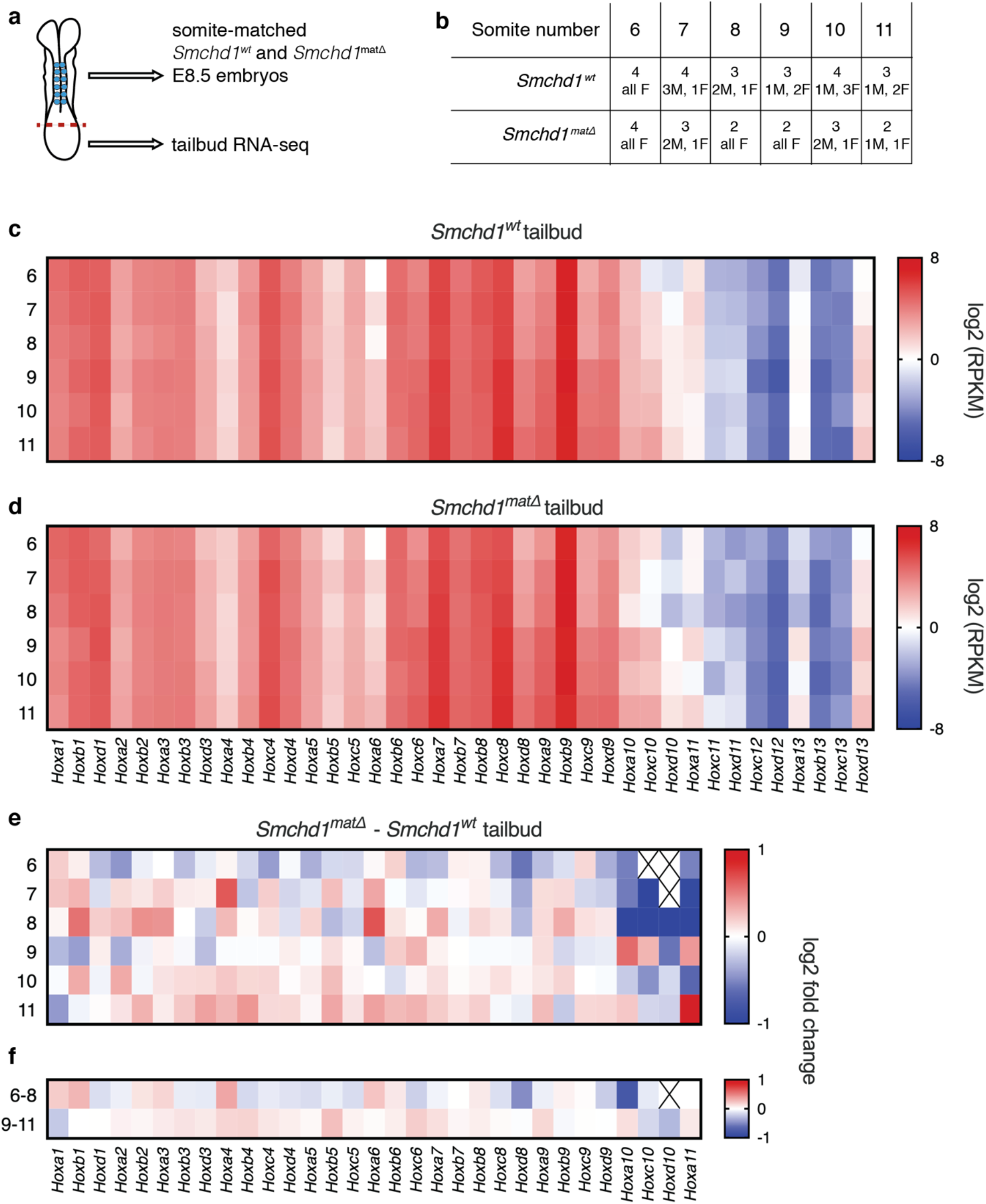
Precocious activation of anterior *Hox* genes in *Smchd1* maternal null tailbud samples. **(a**) Graphical depiction of an ~E8 embryo, showing somites in blue and dissection point by a dotted red line, and a summary of the experimental approach. (**b**) Tabular summary of replicate and sex data for each embryo at each somite stage dissected. (**c-d**) Heatmap of the average reads per million normalised to gene size (RPKM) for the *Hox* genes at 6 to 11 somites in the control (**c**) and *Smchd1* maternally deleted (**d**) tailbud samples. The colour scheme is given. (**e**) Heatmap showing average log2 fold change for expressed *Hox* genes between the *Smchd1* maternal null and control samples. Genes below the expression threshold are denoted with a cross. (**f**) As in (e) but for grouped somite numbers 6-8, 9-11.

To further explore the precocious anterior *Hox* activation, we opted for an *in vitro* approach by differentiating murine embryonic stem cells (mESCs) into NMPs. We derived *Smchd1^wt^* and *Smchd1^matΔ^* mESCs, deleting in the oocyte with Zp3-Cre and performed RNA-seq in the mESCs, where we observed no significantly differentially expressed genes (n=3, Supplementary Dataset 3, Supplementary Figure 2). Next, we differentiated the *Smchd1^wt^* and *Smchd1^matΔ^* mESCs, harvesting RNA from differentiating cells every 12 to 24 hours from their pluripotent to NMP-like state (n=4, Fig. 3a). Just as observed *in vivo*, there was no differential expression when analysed genome-wide (Supplementary Figure 2). The differentiation progressed as expected with the loss of pluripotency factor expression and increase in differentiation factors (Supplementary Figure 2). 12 hours after Wnt activation (day 2.5), we observed precocious activation of several anterior *Hox* genes in the *Smchd1^matΔ^* cells (Fig. 3b), which corresponds to approximately E8.5 *in vivo* ^40^. Given the *Hox* genes are just switching on at this time, this was most noticeable as a larger log fold change between day 2 and day 2.5 of differentiation in the maternal null compared with control cells for anterior *Hox* genes (Fig. 3c, d, p<0.001). At day 3 (NMPs and MPs) and day 4 (24h after GDF11 addition) corresponding to E9.5 *in vivo*, we observed a general downregulation of *Hox* gene expression, consistent with the *Hox* gene activation we observe at day 2.5 being precocious but not sustained (Fig. 3b).

**Figure 3.**
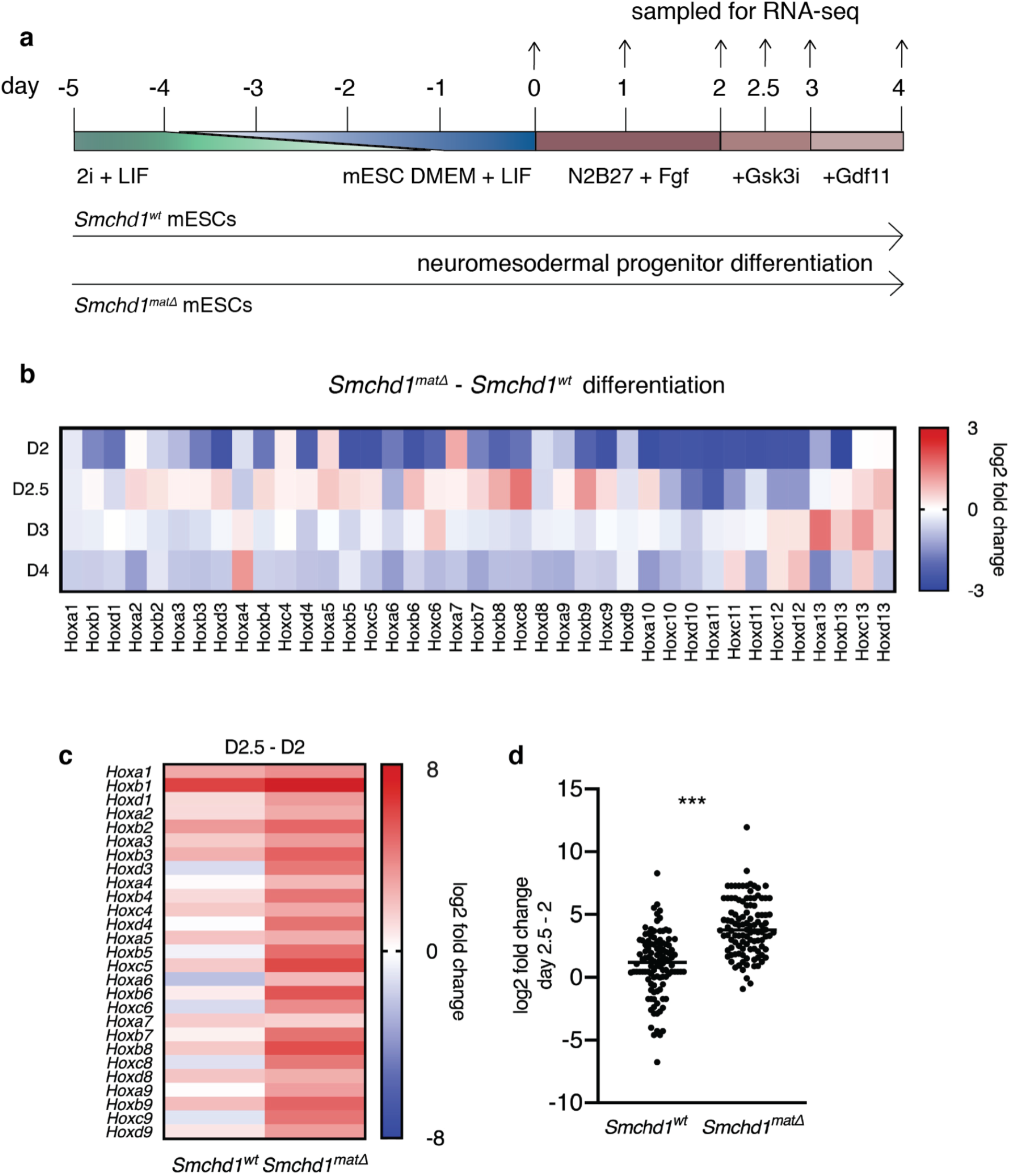
Precocious activation of anterior *Hox* genes in differentiating *Smchd1* maternal null mESCs upon Wnt activation. **a**. mESC differentiation to NMP experimental setup, with media components indicated, along with timing of samples taken for RNA-seq. **b**. Heatmap of the average log2 fold change for the expressed *Hox* genes between the *Smchd1* maternal null and control samples. n=4 replicates for each genotype of mESC, from two separate mESC lines for each genotype. **c**. Heatmap of the average log2 fold change of *Hox* gene expression between the day 2.5 and day 2 samples, for each genotype. **d**. The log2 fold change for the *Hox1-9* genes between day 2.5 and day 2 of differentiation, for each of the 4 replicates per genotype (Student’s t-test, two-tailed, equal variance *** p<0.001).

Taken together, these *in vivo* and *in vitro* data suggest that the *Smchd1^matΔ^* skeletal phenotype may in part be explained by premature upregulation of anterior *Hox* genes. The normal *Hox* gene silencing in the pluripotent state, and the expected downregulation of *Hox* genes observed later in differentiation, is consistent with the relatively modest effects on skeletal patterning in the absence of maternal SMCHD1.

### Maternal SMCHD1 acts downstream of Polycomb-mediated H3K27me3 and H2AK119ub in mESCs

To investigate the mechanism by which loss of maternal SMCHD1 caused upregulation of anterior *Hox* genes, we assessed whether the histone marks H2AK119ub and H3K27me3 were perturbed in *Smchd1^matΔ^* mESCs. These histone marks are laid down by polycomb repressive complex 1 (PRC1) and 2 (PRC2) respectively and induce a heterochromatic gene-silencing state at the *Hox* clusters in mESCs ^41, 42^. Moreover, both marks are laid down on the maternal chromatin at *Hox* clusters and elsewhere in the genome ^34, 35, 36^. For non-canonical imprinted genes, maternal H3K27me3 and H2AK119ub are required for their silent state. Based on recent work from our lab on the role of maternal SMCHD1 at non-canonical imprinted genes ^7^ we hypothesised that maternal SMCHD1 acts downstream of PRC1 and PRC2. If this were the case, H3K27me3 and H2AK119ub would be unperturbed over *Hox* clusters in *Smchd1^matΔ^* mESCs. We carried out CUT&RUN for H3K27me3 and H2AK119ub in biological triplicate samples of *Smchd1^wt^* and *Smchd1^matΔ^* mESCs, and compared our data to publicly available ChIP-seq datasets for these histone marks in mESCs (Supplementary Dataset 4, ^41, 42^). We observed enrichment of H3K27me3 and H2AK119ub at the expected regions for mESCs, including the four *Hox* clusters (Fig. 4a-h); however, we found no change in H3K27me3 or H2AK119ub enrichment between *Smchd1^wt^* and *Smchd1^matΔ^* mESCs over the *Hox* clusters (Fig. 4a-h), and a very high positive correlation between the two genotypes genome-wide (Fig. 4i, j, R^2^=0.9266 and 0.8721, respectively). Moreover, there was no significant difference when considering just the maternal or paternal allele of each cluster (Supplementary Figure 3). Given that the *Smchd1* maternal null mESCs retained their maternal effect on *Hox* gene expression and that loss of maternal SMCHD1 does not disrupt acquisition and/or maintenance of the Polycomb repressive complex marks, maternal SMCHD1 must act downstream of these marks in its regulation of *Hox* clusters and other areas of the genome.

**Figure 4.**
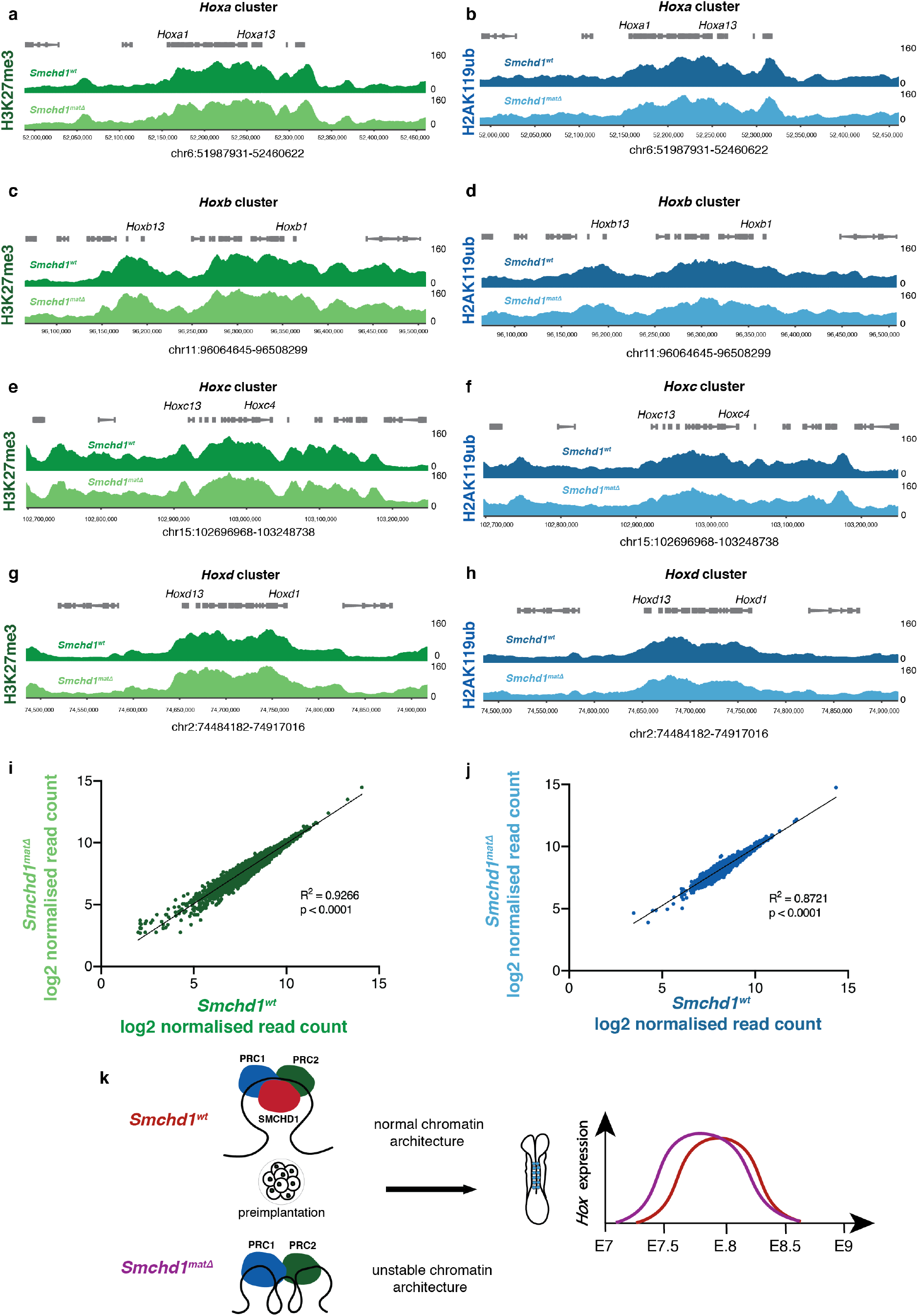
Maternal SMCHD1 acts downstream of H3K27me3 and H2AK119ub in mESCs. **a, c, e, g**. H3K27me3 CUT&RUN in *Smchd1^wt^* and *Smchd1^matΔ^* ESCs cultured in 2i+LIF medium over the four *Hox* clusters as marked. n=3 independent mESC lines per genotype, the average of which is shown. Genes are shown in grey above the CUT&RUN enrichment tracks (log2 FPKM), genome coordinates are shown below. Green indicates H3K27me3, dark green for *Smchd1^wt^* and light green for *Smchd1^matΔ^*. **b, d, f, h**. as in (a) but for H2AK119ub. Blue represents H2AK119ub, dark blue for *Smchd1^wt^* and light blue for *Smchd1^matΔ^*. **i**. Scatter plot of log2 transformed normalised counts for *Smchd1^wt^* and *Smchd1^matΔ^* over H3K27me3 MACS2 peaks in mESCs called from ^41^. The Pearson coefficient indicates very high correlation between the two genotypes. **j**. As in b. but for H2AKA119ub, peaks called from ^42^. Pearson coefficient again indicates very high correlation between the genotypes. **k**. Cartoon illustrating our proposed model that maternal SMCHD1 acts downstream of Polycomb to maintain a stable chromatin architecture at *Hox* genes for normal expression post-implantation.

## Discussion

In this study we have shown that maternal SMCHD1 is required for appropriate patterning of the axial skeleton, linked to its role in silencing *Hox* genes. Deletion of *Smchd1* in the oocyte results in the highly penetrant posteriorising homeotic transformations of C7-to-T1, T13-to-L1 and L6-to-S1. This phenotype is similar to what is observed in zygotic PRC1 subunit knockouts and is attributable to *Hox* gene overexpression ^43, 44^. We too observed a modest but consistent upregulation of anterior *Hox* genes both *in vivo* in the developing tailbud of ~E8.5 embryos and *in vitro* soon after induction of Wnt signalling in mESC differentiating into neuromesodermal progenitors. *Hox* gene silencing was restored later in differentiation, consistent with the relatively subtle axial patterning defects observed in the *Smchd1* maternal null embryos. Interestingly, maternal SMCHD1 was not required to maintain appropriate *Hox* gene silencing earlier in development, either in the pluripotent state or in the morula. These data suggest that maternal SMCHD1, which controls the embryo in the preimplantation period ^7^, is required to ensure *Hox* genes are not prematurely activated in the post-implantation period. This is a long-lived effect of maternal SMCHD1 at the *Hox* clusters from around E2.75 when zygotic SMCHD1 is activated to around E8.5 when precocious *Hox* gene activation is observed.

Given the important role of PRC1 and PRC2 in silencing the *Hox* genes, their role in long-lived mitotic epigenetic memory ^45^ (and the fact that they mark the *Hox* genes with H2AK119ub and H3K27me3 from the oocyte stage onwards ^4, 34, 35, 36, 41^, we asked whether maternal SMCHD1 may function together with the PRCs to silence *Hox* genes, using our mESC model. In undifferentiated mESCs we observed no change in H3K27me3 or H2AK119ub marks genome-wide in *Smchd1* maternally deleted cells compared to control. These data suggest that maternal SMCHD1 acts downstream of polycomb marks in this system, consistent with previous work showing that zygotic SMCHD1 acts downstream of H2AK119ub on the inactive X chromosome ^24^, that H3K27me3 is unchanged over SMCHD1 targets in zygotic *Smchd1* null NSCs ^11^, and the role for maternal SMCHD1 at non-canonical imprinted genes controlled by H3K27me3 and H2AK119ub ^7^. Potentially, the retention of H2AK119ub and H3K27me3 in the absence of maternal SMCHD1 explains why *Hox* genes are not aberrantly expressed prior to E8.5.

In our embryos we observe a fairly restricted set of posterior homeotic transformations, with the highest penetrance observed for transformations at the anterior end of the skeleton and upregulation of anterior *Hox* genes. Recent studies have reported the dynamic changes in H3K27me3 and H2AK119ub through the preimplantation period, relevant to this consideration ^34, 35, 36^. Zheng et al. showed that although H3K27me3 decorates the *Hox* clusters in the oocyte and sperm, paternal H3K27me3 is erased and is regained post-implantation, while maternal H3K27me3 remains constantly at *Hox* loci. Mei et al. showed that the same is true for H2AK119ub except it is regained on the paternal allele by the early two-cell stage, faster than paternal H3K27me3 is regained. Considering Polycomb coverage over both alleles (not an allele-specific analysis), Chen et al. showed both H3K27me3 and H2AK119ub were unchanged from the oocyte to morula stage for the posterior 3’ end of the *Hoxc* cluster. Meanwhile, H3K27me3 was erased over the anterior end of the cluster, from the 1-cell to morula stages. H2AK119ub coverage was also reduced, but only at the 1-cell stage. Seeing as this is the window of time when exclusively maternal SMCHD1 protein is present in the embryo ^7^, maternal SMCHD1 may affect anterior *Hox* genes because lower levels of H3K27me3 and H2AK119ub create a higher dependence on maternal SMCHD1 for an appropriate chromatin state at the *Hox* genes. These studies also show that neither the dynamic changes in Polycomb marks as the early embryo develops, nor the allele-specificity of them, is captured in mESC, as we also find in our CUT&RUN data from mESCs. Hence, although our *Smchd1* maternal null mESCs appear to retain the maternal effect of SMCHD1 as they exhibit anterior *Hox* upregulation, in the future it will be important to study the effects of maternal SMCHD1 in the preimplantation period to fully elucidate the role of maternal SMCHD1 at the *Hox* clusters.

Maternal SMCHD1 acting downstream of Polycomb does not fully answer the question of how deletion of *Smchd1* in the oocyte has such a long-lasting effect on the embryo, days after activation of zygotic SMCHD1. Given that zygotic SMCHD1 has a role in maintaining chromatin architecture, specifically long-range chromatin interactions ^12, 21, 23^, including at *Hox* clusters ^12^, it is possible that this long-lasting epigenetic memory exists in the form of a particular chromatin conformation at *Hox* clusters that is put in place early in development by maternal SMCHD1. Without maternal SMCHD1, we propose that the chromatin state of the *Hox* clusters is destabilised leaving *Hox* genes prone to inappropriate activation over time (Fig. 4k). While the Polycomb marks remain, zygotic SMCHD1 activated at the late morula stage appears to be insufficient to ensure appropriate *Hox* gene silencing later in development.

Potentially this is because zygotic SMCHD1 cannot restore the chromatin architecture required for *Hox* silencing at the late morula stage or afterwards, as the establishment of such a chromatin state needs to occur within the context of the dynamic epigenetic reprogramming that happens earlier in preimplantation development. If the Polycomb marks are sufficient for silencing in the short term, why would a maternal SMCHD1-mediated chromatin state be required to prevent premature *Hox* gene activation post-implantation? The early post-implantation period is another time of wholesale epigenome remodelling as the embryo undergoes germ-layer specification and gastrulation. Potentially a destabilised chromatin state created by the absence of maternal SMCHD1 is liable to disruption in the context of such genome-wide remodelling.

Although further work is required to elucidate how maternal SMCHD1 has a long-lasting epigenetic memory in the developing embryo, this study shows that maternal SMCHD1 is required for appropriate *Hox* gene expression and, consequently, is also required for normal skeletal patterning in the mouse embryo. This work is relevant to our understanding of how maternal proteins influence offspring phenotypes, and may be relevant to humans considering the pathogenic variants in SMCHD1 observed in several human diseases ^16, 17, 18, 19^.

## Acknowledgements

We thank Jessica Martin (WEHI) for her valuable assistance with mouse lines. We thank Dr Andrew Langedam (Monash) for imaging assistance. This work was supported by grants and fellowships from the Australian National Health and Medical Research Council (GNT1098290 to MEB, fellowship GNT1194345 to MEB). NB was supported by an Australian Research Training Program scholarship. MEB was supported by the Bellberry-Viertel Senior Medical Research fellowship. Additional support was provided by the Victorian State Government Operational Infrastructure Support, Australian National Health and Medical Research Council IRIISS grant (9000653). The Australian Regenerative Medicine Institute is supported by grants from the State Government of Victoria and the Australian Government.

## Author contributions

NB acquired, analysed and interpreted the data and drafted the paper. QG, ATdF and AK analysed and interpreted data. TB and KB acquired data. EM contributed to design of the work, data acquisition, analysis and interpretation. MB conceived and designed the work, analysed and interpreted the data. All authors edited the manuscript.

## Methods

### Mouse strains and genotyping

Mice were bred, housed and maintained in accordance with standard animal husbandry procedures and experiments performed were approved by the WEHI Animal Ethics Committee under the animal ethics numbers 2018.004, 2020.048 and 2020.50. *Smchd1*^del/fl^ mice carrying the MMTV-Cre transgene ^46^, and *Smchd1*^fl/fl^ mice carrying the Zp3 Cre transgene ^47^ were created as previously described ^7^ and maintained on the C57BL/6 background.

In this study three crosses were carried out to assess the effect of deleting Smchd1 in the oocyte on the developing embryo. The control cross involved *Smchd1*^fl/fl^ MMTV-Cre^+/+^ or Zp3-Cre^T/+^ females and Castaneus (Cast) *Smchd1*^+/+^ male mice to generate *Smchd1*^fl/+^ (Smchd1^wt^) embryos. The maternal deletion test cross was carried out with *Smchd1*^del/fl^ MMTV-Cre^T/+^ or *Smchd1*^fl/fl^ Zp3-Cre^T/+^ females and Cast *Smchd1*^+/+^ male mice, to generate *Smchd1*^del/+^ (Smchd1^matΔ^) embryos which developed from *Smchd1* homozygous-null oocytes. The third cross was a reciprocal cross between a Cast *Smchd1*^+/+^ female and a *Smchd1^del/fl^; MMTV-Cre*^*T*/+^ male to generate *Smchd1*^*fl*/+^ and *Smchd1*^*del*/+^ (*Smchd1^het^*) embryos. The reciprocal cross was only conducted with MMTV-Cre, and not Zp3-Cre, with the purpose of controlling for the heterozygosity of *Smchd1^matΔ^* embryos. The F1 nature of the embryos allows for allele-specific genomic analysis due to differential SNPs between the C57BL/6 and Cast genomes.

Genotyping was carried out as previously described for Smchd1 and the X and Y chromosomes ^12^; and for the Cre transgene ^48^.

### Skeletal preparations

Whole-mount skeletal staining was performed on E17.5 embryos as previously described ^25^ (Rigueur and Lyons, 2014. Skin and organs were removed, embryos dehydrated and remaining tissue dissolved in acetone. After staining, skeletons were cleared in KOH, washed through a glycerol/water series and imaged in 100% glycerol. Images were acquired with a Vision Dynamic BK Lab System at the Monash University Paleontology Lab. Images were taken with a Canon 5d MkII with a 100mm Macro lens (focus stop 1:3/1:1). Multiple images were taken to extend the focal depth, and stacked in ZereneStacker using the PMax algorithm. Two people independently scored vertebral formulae of each skeleton, blind to genotype and sex.

### Tailbud dissection

Tailbud dissection and somite counting was performed as previously described ^49^. In brief, embryos were dissected in ice-cold DEPC-treated PBS. Tailbud tissue was horizontally dissected at a distance of 1.5 somites below the last segmented somite to ensure no contaminating somite tissue was included. Tailbud tissue was snap frozen on dry ice and stored at −80°C for later RNA extraction. The yolk sac was used for genotyping. Somites were counted before fixing each embryo in 4% DEPC-treated paraformaldehyde at 4°C overnight. Embryos were washed through a graded methanol/PBT (DEPC-treated PBS with 1% Tween (v/v)) series as previously described (E. McGlinn and J. H. Mansfield, 2011) before brief staining in dilute ethidium bromide solution and imaging under a fluorescence dissection microscope to confirm somite counting.

### mESC derivation and culture

mESCs were derived and cultured as previously described ^12, 50^. Females were superovulated with 5 IU folligon (MSD Animal Health Australia) 2 days before mating and 5 IU chorulon (MSD Animal Health Australia) on the day of mating. E3.5 blastocysts were flushed from the uterine horns of these females with M2 medium (Sigma-Aldrich) and were washed twice in 2i + LIF medium [KnockOut DMEM (Life Technologies), 1x Glutamax (Life Technologies), 1x MEM Non-Essential Amino Acids (Life Technologies), 1 X N2 Supplement (Life Technologies), 1 X B27 Supplement (Life Technologies), 1x Beta-mercaptoethanol (Life Technologies), 100 U/mL Penicillin/100 μg/mL Streptomycin (Life Technologies), 10 μg/mL Piperacillin (Sigma-Aldrich), 10 μg/mL Ciprofloxacin (Sigma-Aldrich), 25 μg/mL Fluconazol (Selleckchem), 1000 U/mL ESGRO Leukemia Inhibitory Factor (Merck), 1 μM StemMACS PD0325901 (Miltenyi Biotech), 3 μM StemMACS CHIR99021 (Mitenyi Biotech)] before each blastocyst was plated in an individual well of a non-tissue culture treated 24-well plate by mouth pipetting. Blastocysts were left for 7 days at 37 °C in a humidified atmosphere with 5% (v/v) carbon dioxide and 5% (v/v) oxygen before outgrowths were picked and washed in trypsin-EDTA for 5 minutes, then washed in mESC wash media [KnockOut DMEM (Life Technologies), 10% KnockOut Serum Replacement (Life Technologies), 100 IU/mL penicillin/100 μg/mL streptomycin (Life Technologies)], then 2i+LIF media. Outgrowths were mechanically disrupted by pipetting in the 2i+ LIF media and then were transferred into a 24-well to be cultured as mESC lines. Cell lines were genotyped to check Smchd1 knockout and only male lines were selected and these lines were grown in non-tissue culture treated plates in suspension in 2i + LIF medium at 37°C with 5% (v/v) carbon dioxide and 5% (v/v) oxygen, and were passaged using Accutase (Sigma-Aldrich) every other day.

### Differentiation of mESCs into NMPs

Performed as previously described ^12^ with some adaptations. mESCs growing in 2i + LIF medium were passaged onto tissue culture plates coated with 0.1 % porcine gelatin (Sigma-Aldrich), in 75% 2i + LIF media and 25% mESC DMEM + LIF media [(high-glucose DMEM, 0.085 mM MEM Non-Essential Amino Acids (Life Technologies), 34 mM NaHCO3, 0.085 mM 2-mercaptoethanol (Life Technologies), 100 μg/ml streptomycin, 100 IU/ml penicillin, 15% FBS (Life Technologies), 1,000 U/ml ESGRO leukemia inhibitory factor (Merck), 10 μg/ml piperacillin (Sigma-Aldrich), 10 μg/ml ciprofloxacin (Sigma-Aldrich) and 25 μg/ml fluconazole (Selleckchem)]. 24 hours later, medium was changed to 50% 2i + LIF medium and 25% mESC DMEM + LIF medium, then 25% 2i + LIF medium and 75% mESC DMEM + LIF medium 48 hours later. The following day (day −1 of differentiation) cells were then split using Accutase (Sigma-Aldrich) and were seeded on 6-well plates and on 13 mm circular glass coverslips (Hecht, cat no. 6.071 724) in 12-well plates coated with 0.1 % porcine gelatin (Sigma-Aldrich) at densities of 6.25 × 10^4^ cells/cm^2^ for cells to be harvested at days 0 and 1 of differentiation; and 3 × 10^4^ cells/cm^2^ for cells to be harvested on days 2, 2.5, 3 and 4. 24 hours later (day 0 of differentiation) cells were washed with PBS and the medium was changed to N2B27 medium [1:1 mix of Advanced DMEM/F12 (Life Technologies) and Neurobasal medium (Life Technologies), 0.5× N2 supplement (Life Technologies), 0.5× B27 supplement, 1× Glutamax (Life Technologies), 40 μg/ml BSA Fraction V (Life Technologies), 1× 2-mercaptoethanol (Life Technologies), 100 U/ml penicillin/100μg/ml streptomycin (Life Technologies), 10 μg/ml piperacillin (Sigma-Aldrich), 10 μg/ml ciprofloxacin (Sigma-Aldrich) and 25 μg/ml fluconazole (Selleckchem) supplemented with 10 ng/ml recombinant human basic FGF (Peprotech)]. On day 2 of differentiation, the medium was changed to N2B27 medium with 10 ng/ml recombinant human basic FGF (Peprotech) and 5 μM StemMACS CHIR99021 (Mitenyi Biotech). On day 3 of differentiation, the medium was changed to N2B27 medium with 10 ng/ml recombinant human basic FGF (Peprotech) and 5 μM StemMACS CHIR99021 (Mitenyi Biotech), and 50 ng/ml recombinant GDF11 (Mitenyi Biotech). Cells were harvested for RNA on days 0, 1, 2, 2.5, 3 and 4 by lysing cells in the 6-well plate in RNA lysis buffer (Zymo) and freezing at −80 °C until extraction. Cells grown on coverslips were fixed for immunofluorescence on days 0, 1, 2, 2.5, 3 and 4 as described in the immunofluorescence methods section.

### Immunofluorescence

Immunofluorescence was performed on differentiating mESCs to NMPs as previously described ^12^. In brief, cells grown on coverslips were 13 mm circular glass coverslips (Hecht, cat no. 6.071 724) were washed 3 times for 5 minutes each in PBS before fixation in 4% paraformaldehyde for exactly 10 minutes. Fixed cells were then stored in 0.02% sodium azide in PBS at 4 °C for up to a week until all samples from differentiation were ready for processing. Cells were then washed 3 times for 5 minutes each again in PBS before permeabilisation in 0.5% TritonX in PBS for exactly 5 minutes on ice. Cells were washed again 3 times for 5 minutes each in PBS then non-specific binding sites were blocked in 1% bovine serum albumin (Sigma-Aldrich, A9418) in PBS for approximately 1 hour at room temperature. Primary antibodies against T/Brachyury (Abcam, #ab209665) and Sox2 (ThermoFisher Scientific, cat #14-9811-82) were then added in 1% BSA solution at a dilution of 1:100 and incubated with cells at 4 °C overnight in a humidified chamber. Cells were then washed 3 times for 5 minutes each in PBS before incubation with secondary goat anti-rabbit 647 (ThermoFisher Scientific, cat #A21244) and goat anti-rat 568 (Invitrogen, cat #A11077) antibodies in a dark humidified chamber for one hour at room temperature, before washing 3 times for 5 minutes each again in PBS and counterstaining with DAPI 1:10,000 in PBS for 1 minute at room temperature. Cells were washed 3 times for 5 minutes each again in PBS before mounting on Polysine microscope slides (LabServ, cat #LBSP4981) with Vectashield Vibrance Antifade mounting medium (Vector Laboratories, cat #H-1700). Cells were imaged on an LSM 880 (Zeiss) confocal microscope at 40X magnification and z-stacks were merged and composite images generated using the ImageJ distribution package FIJI ^51^.

### CUT&RUN

CUT&RUN was performed as previously described ^52^. mESCs grown in 2i media were taken out of culture and counted using a haemocytometer before washing by centrifuging at 600 g for 5 minutes and room temperature then resuspending in wash buffer [20 mM HEPES pH 7.5, 150 mM NaCl, 0.5 mM spermidine and 1X complete protease inhibitor (Roche)] three times. Approximately 200,000 cells were used for each antibody for replicate 1, and 500,000 each for replicates 2 and 3. 10 uL Concanavalin A-coated beads (Bangs Laboratories, #BP531) per sample were washed in binding buffer (20 mM HEPES pH 7.5, 10 mM KCl, 1mM CaCl2, 1mM MnCl2) then were resuspended in the original volume of beads. 10 uL beads per sample were then bound to the cells in 1 mL wash buffer, nutating for 10 minutes at room temperature. H3K27me3 (Cell Signalling Technologies, C36B11) and H2AK119ub (Cell Signalling Technologies, D27C40) antibodies were added at a concentration of 1:100 in antibody binding buffer (2 mM EDTA in digitonin wash buffer (wash buffer with 0.025% digitonin) and antibody binding was conducted overnight at 4°C with both H3K27me3, rotating on a nutator. Samples were then washed three times with digitonin wash buffer before resuspending in digitonin wash buffer with pAG-MNase (EpiCypher, 15-1116) at a concentration of 1:20 and nutating at 4°C for 1 hour to allow pAG-MNase and epitope binding. Samples were then washed twice with digitonin wash buffer then once with low-salt rinse buffer (20 mM HEPES pH 7.5, 0.5 mM spermidine, 0.05% digitonin) before resuspending in 200 uL ice cold incubation buffer (3.5 mM HEPES pH 7.5, 10 mM CaCl2, 0.05% digitonin). Samples were then incubated at 0 °C for exactly 30 minutes to allow MNase cleavage at antibody bound sites, before resuspension in 200 uL STOP buffer (170 mM NaCl, 20 mM EGTA, 0.05% digitonin, 50 ug/mL RNase A, 25 ug/mL glycogen) then incubation at 37 °C for 30 minutes. Supernatant containing the cleaved chromatin was separated from the ConA beads, DNA was extracted and purified using phenol/chloroform/isoamylalcohol followed by ethanol and glycogen precipitation, purified DNA was resuspended in 0.1% TE buffer (1 mM Tris-HCl pH 8.0, 0.1 mM EDTA). DNA was quantified with a Qubit dsDNA HS Assay Kit and 6ng or total DNA if less than 6ng was used as the input for sequencing library preparation.

Libraries were prepared using the NEBNext Ultra II DNA Library Prep Kit for Illumina (New England Biolabs, E7645), with adapters diluted 1:5 from the supplied concentration. Libraries were quantified on a HS D1000 tape on a 4200 Tapestation (Agilent Technologies) before pooling and sequencing on the Illumina NextSeq platform, with 75 bp paired-end reads.

### CUT&RUN analysis

Adapter trimming of Fastq files was performed using TimGalore! v0.4.4 with Cutadapt v1.15 ^53^ then QC was carried out using FastQC v0.11.8. Reads were mapped to the GRCm38.p6 version of the mouse reference genome using Bowtie2 v2.3.4.1 ^54^ and Samtools v1.7 ^55^. For allele-specific analysis, reads were mapped to the GRCm38 mouse genome reference N-masked for Cast SNPs prepared with SNPsplit v0.3.2 ^56^. Bam files were imported into SeqMonk v1.47.1, v1.47.2 or v1.48.0 ^57^. H3K27me3 and H2AK119ub MACS peaks were called from publicly available ChIP-sequencing datasets (settings for 300 bp, p < 1 × 10-5) (^41^ GEO accession number GSE73952, ^42^ GEO accession number GSE161996) between the histone mark ChIP and input libraries. The three biological replicates each for *Smchd1^wt^* and *Smchd1^matΔ^* were merged, and peaks were called against the MACS peaks from the publicly available data using the feature probe generator function in SeqMonk (±2.5kb). Probes were quantitated using the read based quantitation option, normalising to library size and log2 transforming the read count. Read counts were exported from SeqMonk and correlation scatterplots were made using GraphPad Prism v9.0.0. CUT&RUN browser tracks were made by quantifying probes over 1000 bp windows (normalising to library size), sliding by 500 bp, smoothing peaks over 20 adjacent probes.

### RNA-sequencing

RNA-sequencing was carried out on tailbud tissue, 2i mESCs and differentiating cells by first extracting RNA using a Quick-RNA Miniprep Kit (Zymo) with DNase I treatment according to the manufacturer’s instructions. 100 ng total RNA (or less if <100 ng was yielded from a PSM) was used to prepare libraries using either a TruSeq RNA Library Prep Kit v2 (Illumina) or a TruSeq Stranded mRNA kit (Illumina) according to the manufacturer’s instructions. Libraries were size-selected for 200-600 bp and primer dimers cleaned up using Ampure XP beads (Beckman Coulter Life Sciences) and were quantified using a D1000 tape on a 4200 Tapestation (Agilent Technologies). Libraries were then pooled and sequenced on the Illumina NextSeq platform, with 75 bp single-end reads, except for 2i mESC RNA-seq libraries which had paired-end sequencing.

### RNA-sequencing analysis

For tailbud, mESC and NMP differentiation RNA-seq, adapter trimming of Fastq files was performed using TimGalore! v0.4.4 with Cutadapt v1.15 ^53^ then QC was carried out using FastQC v0.11.8. Reads were mapped to the GRCm38.p6 version of the mouse reference genome using hisat2 v2.0.5 ^58^ and Samtools v1.7 ^55^. Bam files were imported into SeqMonk v1.47.1, v1.47.2 or v1.48.0 ^57^. Libraries were quantified using Seqmonk’s RNA-seq quantitation pipeline, correcting for total library size and transcript length to generate log2 RPKM values. Differences in *Hox* gene expression were quantified by subtracting *Smchd1^wt^* from *Smchd1^matΔ^* log2 RPKM counts. Heatmaps were generated using GraphPad Prism v9.0.0. Differential gene expression analysis was carried out using the inbuilt edgeR analysis package ^59, 60^ SeqMonk.

For E2.5 male embryo RNA-seq, data was obtained from and analysed as per ^7^. Read were trimmed using TrimGalore v0.4.4 and mapped using hisat2 v2.0.5 to an N-masked version the GRCm38 mouse reference genome for Cast SNPS, made with SNPsplit v0.3.2 ^56^, in paired-end mode and disabling soft-clipping. Gene counts were obtained from bam files in R v3.5.1 ^61^ with the featureCounts function from the Rsubread package v1.32.1 ^62, 63^ provided with the GRCm38.90 GTF annotation downloaded from Ensembl, ignoring multi-mapping or multi-overlapping reads. Gene counts were normalized in edgeR v3.24.0 ^59, 60^ with the TMM method ^64^. Differential gene expression between the Smchd1 maternally deleted and wildtype embryos was performed using the glmFit and glmLRT functions. P-values were corrected with the Benjamini-Hochberg method ^65^. Differential expression results were visualized with Glimma 2.2.0 ^66, 67^.

All genomic data is available on the Gene Expression Omnibus under number GSE183740.

## Supplementary Data

**Supplementary Dataset 1**. Full skeletal scoring data, including comparison of skeletal phenotype in males and females.

**Supplementary Dataset 2.** Raw read count over all genes, Log2 RPKM read counts for *Hox* genes, differential gene expression summary for tailbud samples for merged replicates, and replicates separately.

**Supplementary Dataset 3.** Raw read count over all genes, Log2 RPKM read counts for *Hox* genes and pluripotency/differentiation factors for merged replicates, and replicates separately.

**Supplementary Dataset 4.** Raw read count over all genes, Log2 normalised reads over MACS2 peaks from published datasets for merged CUT&RUN replicates, and replicates separately.

**Supplementary Figure 1.**
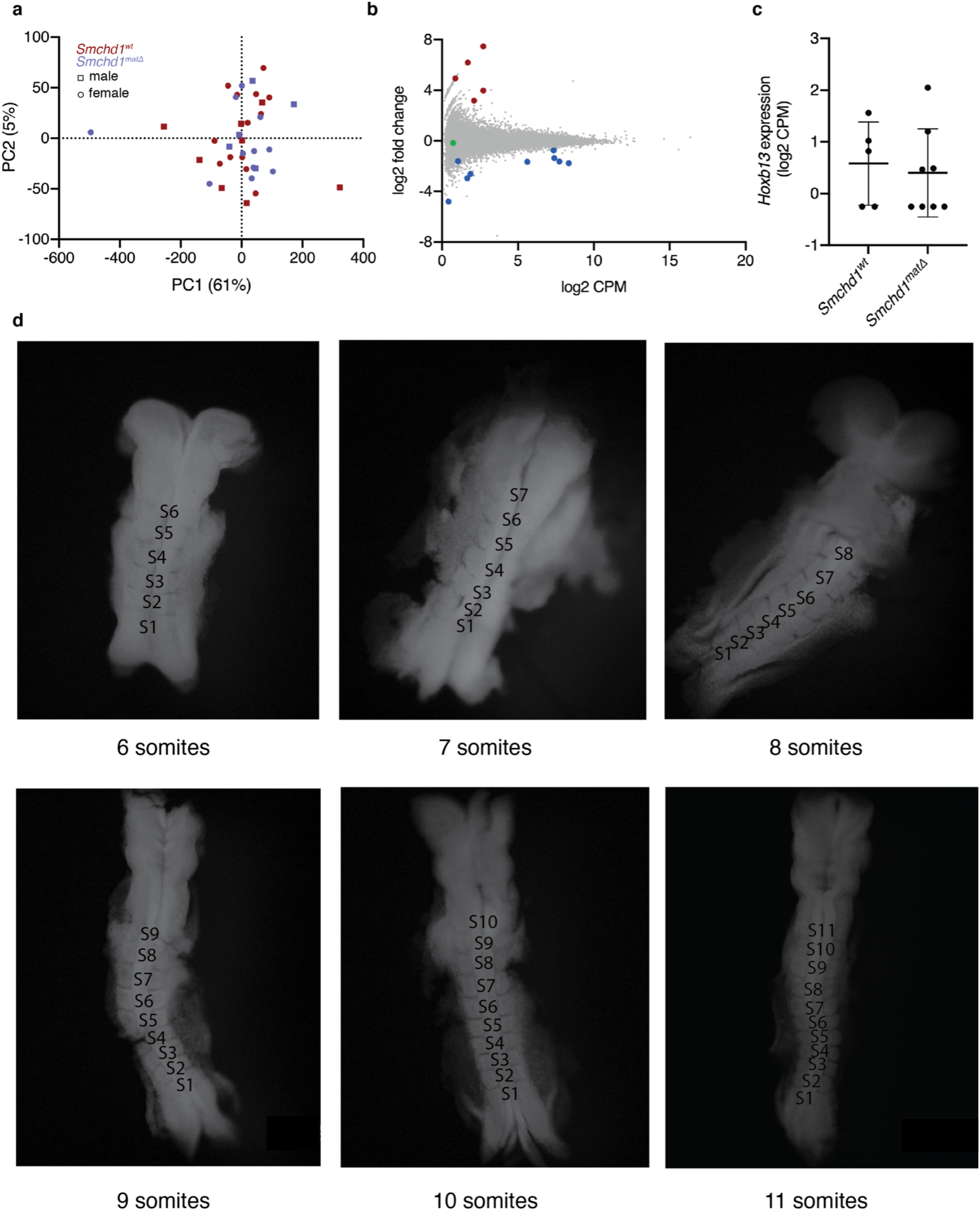
RNA-seq expression analysis in embryos. **a**. Principal Component analysis (PCA) of all RNA-seq samples from E8.0-8.5 tailbud. Red squares represent control *Smchd1*^*fl*/+^ (*Smchd1^wt^*) males, red circles represent *Smchd1^wt^* females, purple squares represent *Smchd1^matΔ^* males, purple circles represent *Smchd1^matΔ^* females. **b**. MA plot of differential expression in *Smchd1^matΔ^* males compared with *Smchd1^wt^* E2.75 embryos, from data published in ^7^. The x-axis shows log counts per million (CPM), while the y-axis is the log2 fold change between the *Smchd1^matΔ^* and control embryos. Significantly differentially expressed genes are shown in red (upregulated) and blue (downregulated). *Hoxb13* is the only detectable *Hox* gene and is shown in green. **c**. *Hoxb13* expression in logCPM in E2.75 male embryos from (b), mean plotted with SD. **d.** Representative dorsal-view images of E8.0-8.5 embryos at each somite stage after tailbud dissection. Note the ventral view of the 8 somite embryo is shown.

**Supplementary Figure 2.**
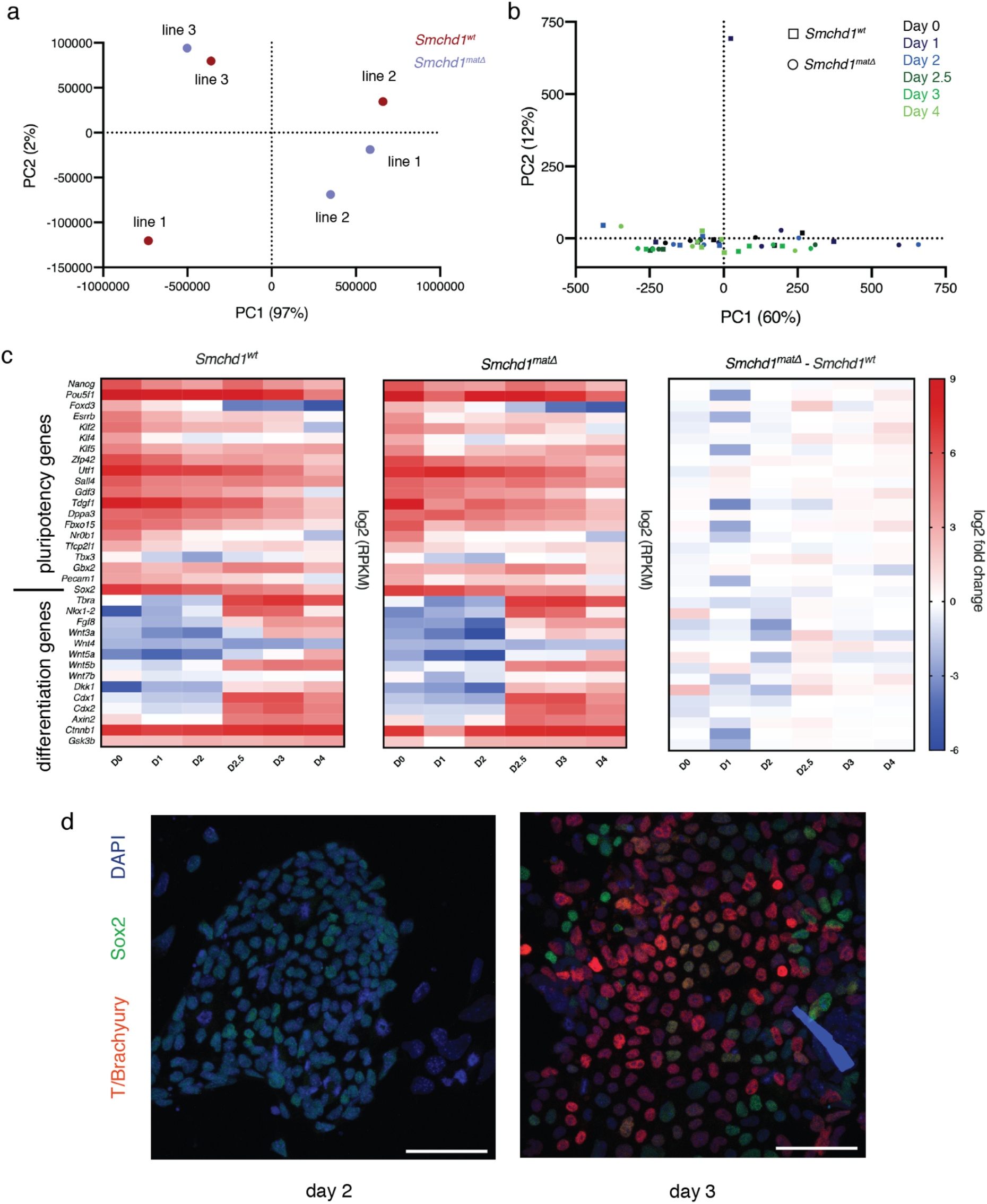
Expression analysis for differentiating *Smchd1^wt^* and *Smchd1^matΔ^* male mESC. **a.** PCA plot for RNA-seq performed on *Smchd1*^*fl*/+^ (*Smchd1^wt^*) and *Smchd1^matΔ^* male mESC maintained in 2i+LIF medium. n=3 independent mESC lines per genotype. **b**. PCA plot for RNA-seq timecourse from day 0 to day 4 of differentiation as shown in Figure 3a. n=3-4 per genotype, representing 2 independent mESC lines per genotype performed 1-2 times each. **c**. Heatmaps of average RPKM of differentiation and pluripotency factor gene expression, in *Smchd1^wt^* and *Smchd1^matΔ^* samples, and the average log2 fold change between these two genotypes. Differentiation and pluripotency factor list compiled from (Gouti et al., 2017, Gouti et al., 2014, Keniry et al., 2019) **d**. Representative IF images of Sox2 and T/Brachyury at days 2 and 3 of differentiation. Scale bar = 50 μm.

**Supplementary Figure 3.**
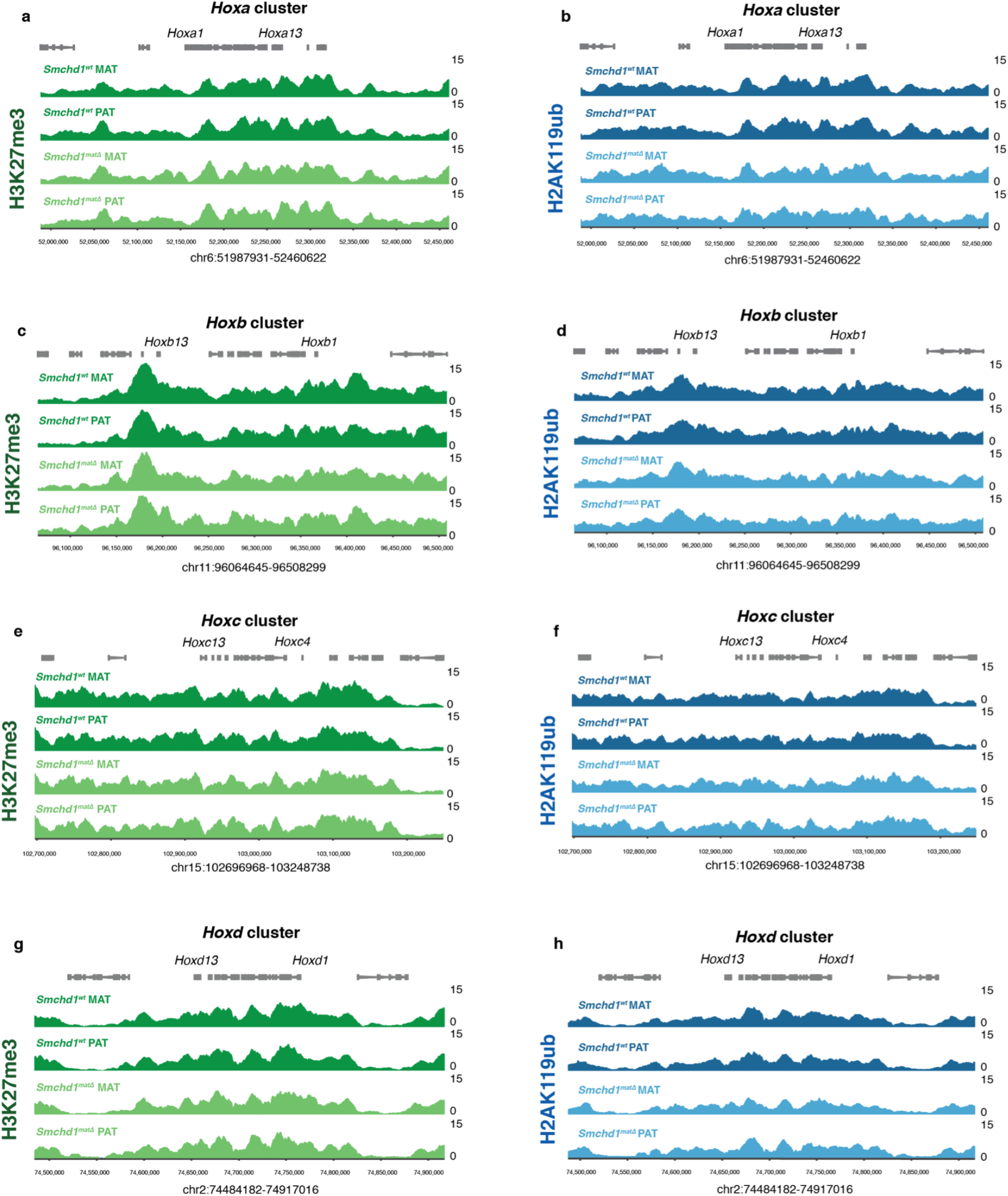
H3K27me3 and H2AK119ub over the maternal and paternal allele of each *Hox* cluster. **a, c, e, g.** CUT&RUN analysis of H3K27me3 over each *Hox* cluster in *Smchd1^wt^* and *Smchd1^matΔ^* ESCs cultured in 2i+LIF medium, where the reads have been split based on SNPs between the C57BL/6 maternal (MAT) and Cast paternal (PAT) genomes. n=3 independent mESC lines per genotype, the average of which is shown. Genes are shown in grey above the CUT&RUN enrichment tracks, genome coordinates are shown below. Green indicates H3K27me3, dark green for *Smchd1^wt^* and light green for *Smchd1^matΔ^*. Note the read counts are significantly lower than for Figure 4 as reads without an informative SNP are removed. **b, d, f, h**. as in a. but for H2AK119ub. Blue represents H2AK119ub, dark blue for *Smchd1^wt^* and light blue for *Smchd1^matΔ^*.

